# Emotional Content Enhances Neural Tracking of Conversations

**DOI:** 10.1101/2025.11.06.686308

**Authors:** P. Heikkinen, S. Lepistö, P. Wikman, E. Koskinen, A. Peräkylä, N. Ravaja, V. J. Harjunen

## Abstract

Engagement in interaction arises through the continuous monitoring of conversation content, allowing individuals to detect and act upon information that is personally relevant or socially significant. Emotional salience of speech content is a key factor in guiding this attentional engagement. By combining an audiovisual “Cocktail-party” paradigm with source-localized magnetoencephalography and encoding–decoding models, we examined whether emotionally salient content continuously enhances attentional tracking of conversations. Decoding analyses showed that emotional salience strengthened neural speech tracking across the conversational arc, but only when dialogue was attended to. The emotional enhancement was also sensitive to the dialogue’s semantic structure, peaking earlier in semantically coherent than incoherent conversations. Encoding analyses revealed latency-specific effects across temporal, parietal and frontal regions. Emotional salience and semantic structure co-modulated mid-latency sensory encoding, whereas later semantic integration was driven by emotionality alone. The observed results support the notion of adaptive tuning of auditory–language networks to socially and motivationally significant semantic cues. Rather than acting as a simple amplifier of semantic processing, emotional salience of dialogue topics shapes neural speech processing at multiple stages. The exact modulations depend on the semantic structure of the dialogue, balancing the needs between efficiency and adaptability that together support the ability to engage optimally in real-world interactions.

**Significance statement:** The emotionality of a conversation is a key factor in how people allocate attention during interaction. Using a multi-speaker “cocktail-party” paradigm with magnetoencephalography, we show that emotionally salient content and semantic coherence influence the brain’s tracking of continuous dialogues in the presence of competing speech. Emotionally salient content enhances neural tracking only for attended speech, and this enhancement appears earlier when the conversation consists of semantically interrelated turns and is easier to follow. This is supported by modulations in neural activity at distinct processing stages across a distributed cortical network. These findings indicate that emotionality functions as a relevance signal, sustaining attentional engagement in conversation by affecting predictive processing and modulating activations in a large-scale cortical system linked to speech comprehension.

## Introduction

Staying focused on what’s being said is essential for understanding and contributing meaningfully to a conversation. Emotional content plays a powerful role in this process: cues that carry strong emotional significance and behavioral relevance heighten sensory processing by activating attention-related neural systems (e.g., Lang et al., 1998; Sander et al., 2003; Yiend, 2010). Consider a social gathering, news of an accident or crime will almost inevitably draw more attention than a casual chat about commuting or grocery shopping.

While much neurocognitive research on emotional language processing has emphasized the role of local cues, such as prosodic variation or individual word choices (e.g., Berckmoes & Vingerhoets, 2004; Citron, 2012), meaning in natural language is not conveyed solely by these elements. Instead, semantic meaning unfolds continuously as words and sentences are integratively processed within an evolving contextual structure (Caucheteux, 2023; Xu et al., 2005). Understanding how emotional meaning sustains attentional engagement in speech therefore requires considering neural processing in contexts of naturalistic, continuous speech.

At the neural level, much of what we know about emotional speech/language processing comes from studies examining cortical responses to local emotional cues. Here, the body of evidence shows that emotionality in word meaning and intonation influences multiple stages of speech processing ranging from early sensory encoding (e.g., N1), salience detection (e.g., P2), to later evaluative processing (e.g., N400) (Kotz & Paulmann, 2011; Schirmer & Kotz, 2006; Pinheiro, 2025). However, even the processing of these local emotional speech cues is sensitive to surrounding linguistic context (Bayer et al., 2010; Ding et al., 2015, 2016; Fields & Kuperberg, 2012; Martín-Loeches et al., 2012). For example, emotionally congruent contexts reduce neural response amplitudes to emotional words, mirroring facilitatory effects of semantic predictability (Nieuwland & Van Berkum, 2006; Van Berkum et al., 2003; Zhang et al., 2021). These findings indicate that neural processing of emotional word-meanings depends on the context they appear in.

In natural speech, such as during storytelling or a conversation, social and emotional meanings emerge across extended stretches of discourse rather than being contained in individual word choices (Hoemann et al., 2025; Mar, 2011). We propose that this accumulating emotional salience acts as a high-level relevance signal that continuously biases attentional engagement. Research using the cocktail-party paradigm, with multiple concurrent speech streams, has shown that such attentional engagement is reflected in enhanced neural tracking of the attended stream (Cherry, 1953; Ding & Simon, 2012; Fiedler et al., 2019; O’Sullivan et al., 2015; Zion Golumbic et al., 2013). This facilitates segregating the speakers into distinct neural representations by integrating information across multiple timescales, while biasing resource-allocation for higher-order semantic processing to the attended stream (Brodbeck et al., 2022; Power et al., 2012; Teoh et al., 2022). Recent work further demonstrates that neural tracking of conversational speech fluctuates nonlinearly across dialogue turns and is supported by a highly distributed cortical network (Wikman et al., 2021, 2024).

Building on these findings, we hypothesize that emotional salience enhances neural tracking of conversational speech, especially when it is the focus of attention. We further predict that the narrative structure of the conversations modulates the dynamics of neural tracking, as it breaks down the contextual cues facilitating neural processing. Finally, we hypothesized that emotional salience and narrative structure would jointly modulate distinct stages of neural speech processing, with effects emerging most strongly at stages associated with semantic evaluation in cortical regions implicated in language comprehension (Fedorenko, 2024; Ji et al., 2019).

## Results

Using magnetoencephalography (MEG), we recorded brain activity while participants selectively attended to lifelike audiovisual target conversations and were asked to ignore an overlapping audio-only distractor conversation in a “cocktail party” paradigm (Wikman et al., 2021, 2024). We factorially manipulated both emotional salience of the conversation topic (emotional vs. neutral) and semantic dialogue coherence across turns (coherent vs. incoherent) in both target and distractor conversations (Fig. 1A). To examine how each factor shapes attentional tracking of conversational speech, all factor combinations in target dialogues were presented with all combinations in the distractor dialogues (Fig. 1B). All speech was presented with flat prosody and a neutral facial expression, to ensure the observed effects won’t reflect lower-order paralinguistic and nonverbal emotional cues.

**Figure 1.**
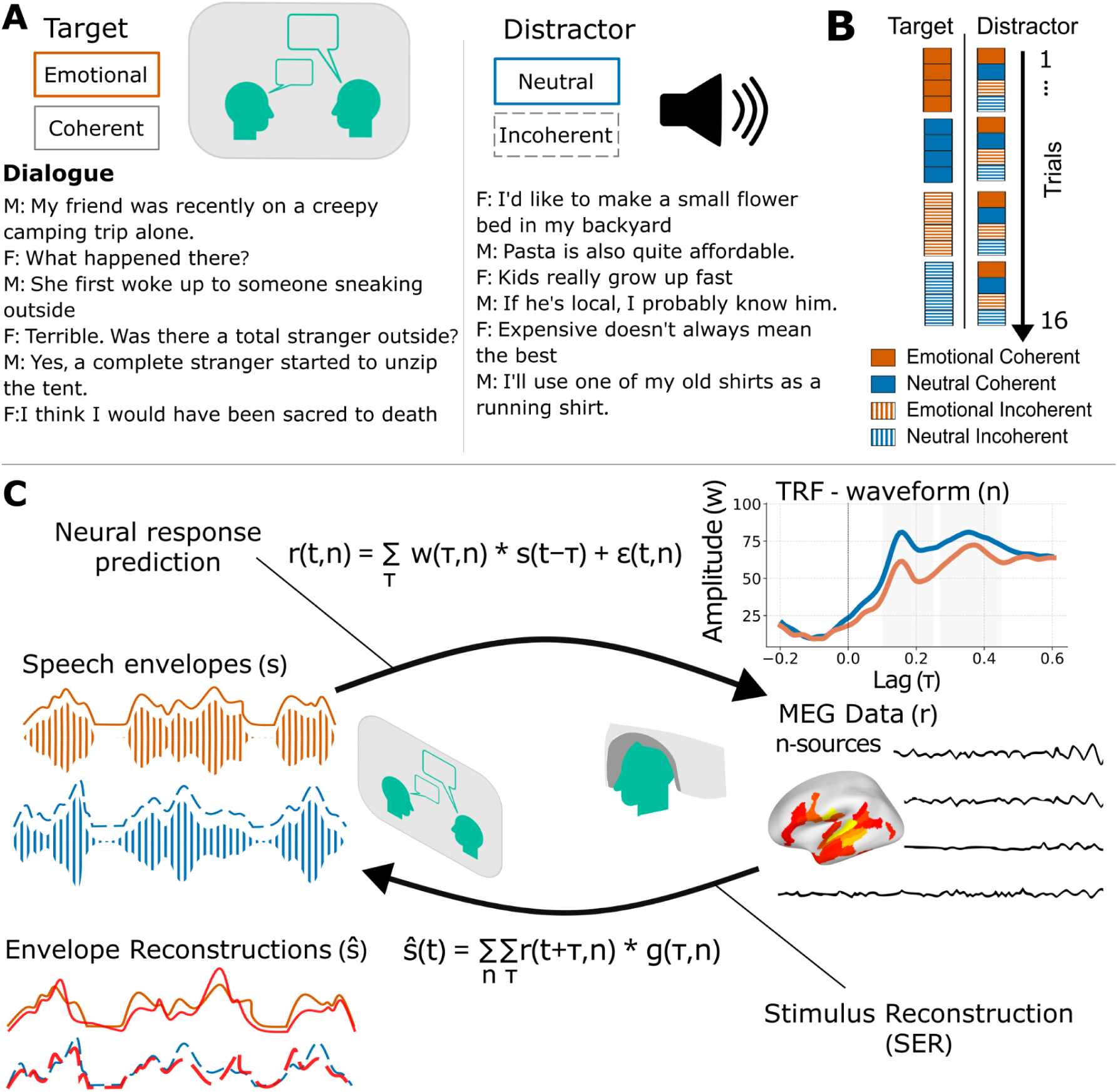
The experimental design and analysis framework. (A) Example stimulus materials. The left panel shows an example of an audiovisual target dialogue, including the video stimulus and transcript, illustrating an emotionally salient coherent dialogue (see SI Methods for details on dialogue content). The right panel shows a corresponding audio-only distractor dialogue, exemplifying a neutral incoherent conversation. (B) Structure of a single trial (one of two per participant), in which each target dialogue type was presented with each distractor type. Trials are shown grouped by target condition for illustration; actual presentation order was pseudorandomized (Methods). (C) Schematic overview of the modelling approaches. Speech envelopes of the target (orange) and distractor (blue) dialogues for each condition were used separately to predict how neural activity responds to changes in the speech envelope with temporal response functions (TRFs). In the speech envelope reconstruction (SER) analysis, neural activity was used to reconstruct the corresponding speech envelopes. Note that presented reconstructions and TRF waveforms are for illustrative purposes only.

Neural responses were analyzed with a combined encoding-decoding approach, utilizing temporal response functions (TRFs) and speech envelope reconstruction (SER; Fig. 1C; Crosse, 2016; Lalor & Foxe, 2010). In SER, the multivariate time series of neural activity serves as input to reconstruct the acoustic speech envelope. Speech-related information is distributed across MEG-electrodes and time points, and the decoding model leverages this multidimensional representation to maximize stimulus recovery. TRFs were used to model how changes in the continuous speech envelope drove cortical activity. TRFs are physiologically interpretable, linear combinations that describe timing and strength of stimulus–response relationships as specific time-lags and are thus analogous to event-related potentials (Crosse et al., 2021; Kaufman & Zion Golumbic, 2023).

### Emotional Content and Dialogue Structure Affect Perception and Recall

Participants rated the perceived emotional load of each dialogue and rated statements about their content either true or false. Regarding emotional load (rated on a three-point scale), emotional dialogues (*M* = 2.40, *SE* = 0.69), in contrast to neutral ones (*M* = 1.06, *SE* = 0.24), were perceived as more affectively charged (ordinal-GEE: Wald χ²(1) = 10,867.51, *p* < .001; SI Table S1). This indicated that participants were sensitive to the emotionality embedded in the conversation content. With respect to the coherence, participants’ recall accuracy (proportion of correct answers to true–false statements) for the coherent dialogue content was higher (*M* = 0.89, *SE* = 0.20) than for incoherent ones (*M* = 0.80, *SE* = 0.21), implying that contextual continuity had facilitatory effect on memory performance (rm-ANOVA: *F*(1, 30) = 53.73, *p* < .001, η*_G_*^2^ = 0.199, SI Table S2). Emotional salience did not affect recall accuracy (*p* = .376, η*_G_*^2^ = 0.004).

### Emotional Content Enhances Neural Tracking of Attended Conversations

SER showed better accuracy for attended, compared to distractor speech, demonstrating a robust attentional enhancement in neural speech tracking (rm-ANOVA: *F*(1, 30) = 408.79, *p* < .001, η*_G_*^2^ = 0.56). SER accuracy was also higher for emotional than for neutral dialogues (*F*(1, 30) = 26.66, *p* < .001, η*_G_*^2^ = 0.06), whereas coherence had no impact on overall tracking accuracy (*p* = .83; SI Table S3). Importantly, the influence of emotional salience depended on attentional engagement: emotional content enhanced SER only for attended speech, regardless of dialogue coherence (Fig. 2A; Attention x Emotional Salience x Coherence interaction: *F*(1, 30) = 6.17, *p* = 0.019, η*_G_*^2^ = 0.02; SI Table S4).

**Figure 2.**
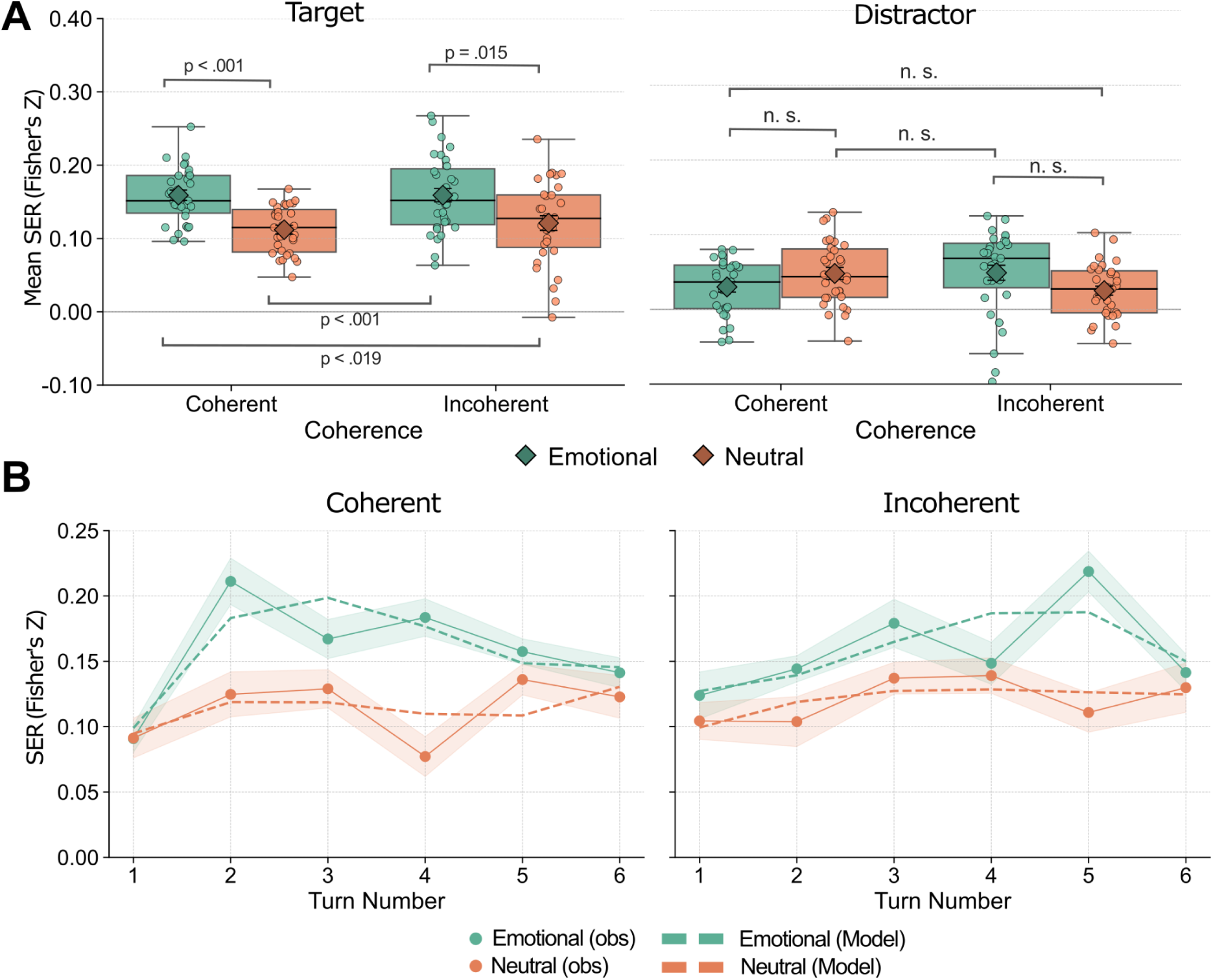
Speech Envelope Reconstruction Results. (A) Mean SER accuracy (Fisher’s z-transformed) for attended and distractor speech as a function of emotional salience and dialogue coherence. Boxplots show empirical distributions, with individual observations overlaid. Diamonds indicate estimated marginal means with standard errors. (B) Turn-by-turn trajectories of SER accuracy for attended dialogues. Points and solid lines represent observed means with standard error bands. Dashed lines show cubic model-predicted trajectories, illustrating a rapid early peak for coherent emotional dialogues and a slower, sustained increase for incoherent emotional dialogues.

As neural tracking of lifelike dialogues has been shown to display variability in the time domain (Wikman et al., 2021, 2024), we further examined how neural tracking unfolds across conversational turns. Emotional salience modulated the temporal evolution of SER and was modulated by dialogue coherence (Fig. 2B; rm-ANOVA: *F*(4.23, 126.98) = 4.58, *p* = 0.001, η*_G_*^2^ = 0.03; SI Table S5). Specifically, Emotional salience selectively increased the parabolic arc of the SER trajectory in coherent dialogues (SI Table S6), whereas dialogue coherence modulated the timing of emotional effects, with coherent emotional dialogues showing an earlier peak, compared to a more sustained increase in incoherent dialogues (SI Table S7).

Together, these patterns show that emotional content influences both the magnitude and the temporal evolution of neural tracking of speech during active listening, with broader contextual structure determining whether emotional effects appear as a rapid early boost or as a slower, sustained increase across turns.

### Neuronal dynamics of Emotional Speech Processing

To examine how emotional salience affects the temporal and topographical organisation of neural speech processing, we analyzed TRFs for emotionally salient and neutral dialogues in both coherent and incoherent conditions. Grand averaged TRFs exhibited a distinct two-peak structure, with the strongest activations in temporal-frontal regions (Fig. 3A, Methods). Based on these patterns, cortical regions associated with speech and narrative comprehension from the lateral surface were chosen for further analysis (Fig. 3B, Methods; Fedorenko et al. 2024; Glasser et al, 2016; Ji et al, 2019; Simony et al. 2016).

**Figure 3.**
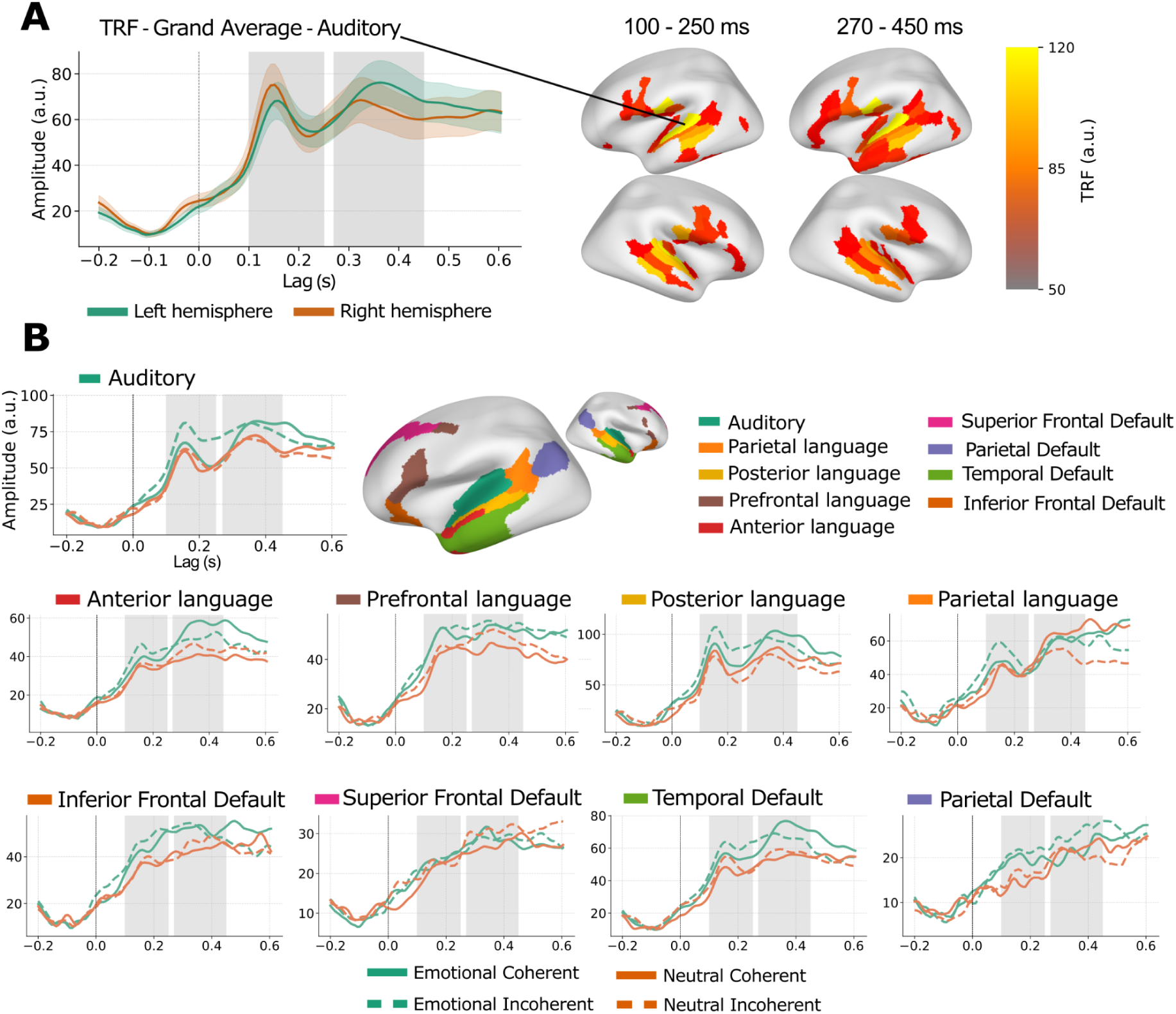
TRF responses and their topographical organization. (A) Grand-averaged TRF waveform in the auditory cortex for both hemispheres across all conditions. The grey highlights denote the mid-latency (100-250 ms) and late-latency (270-450 ms) windows from which mean activations were extracted for statistical analysis. The parcel-level topographies of TRF responses within the chosen time-windows are depicted on the right. (B) Regions of interest (ROIs) chosen for statistical analysis, and left hemisphere TRF-waveforms for all conditions in the target dialogues (see SI Fig. S1, for corresponding right-hemisphere waveforms).

In the mid-latency TRF window (100–250 ms), the effect of emotional salience on neural responses varied across brain areas depending on the narrative structure of the dialogue (Emotion x Coherence x ROI: *F*(4.11, 123.21) = 4.96, *p* < 0.001; Fig. 4A, 4B ; SI Table S8). Emotional salience increased response strength broadly in temporal and parietal regions, while coherent dialogues showed emotional enhancement primarily in the frontal regions (SI Fig. S2). This interaction reflects context-dependent emotional effects that varied across temporal and frontal regions at a latency related to early emotional–semantic evaluation.

**Figure 4.**
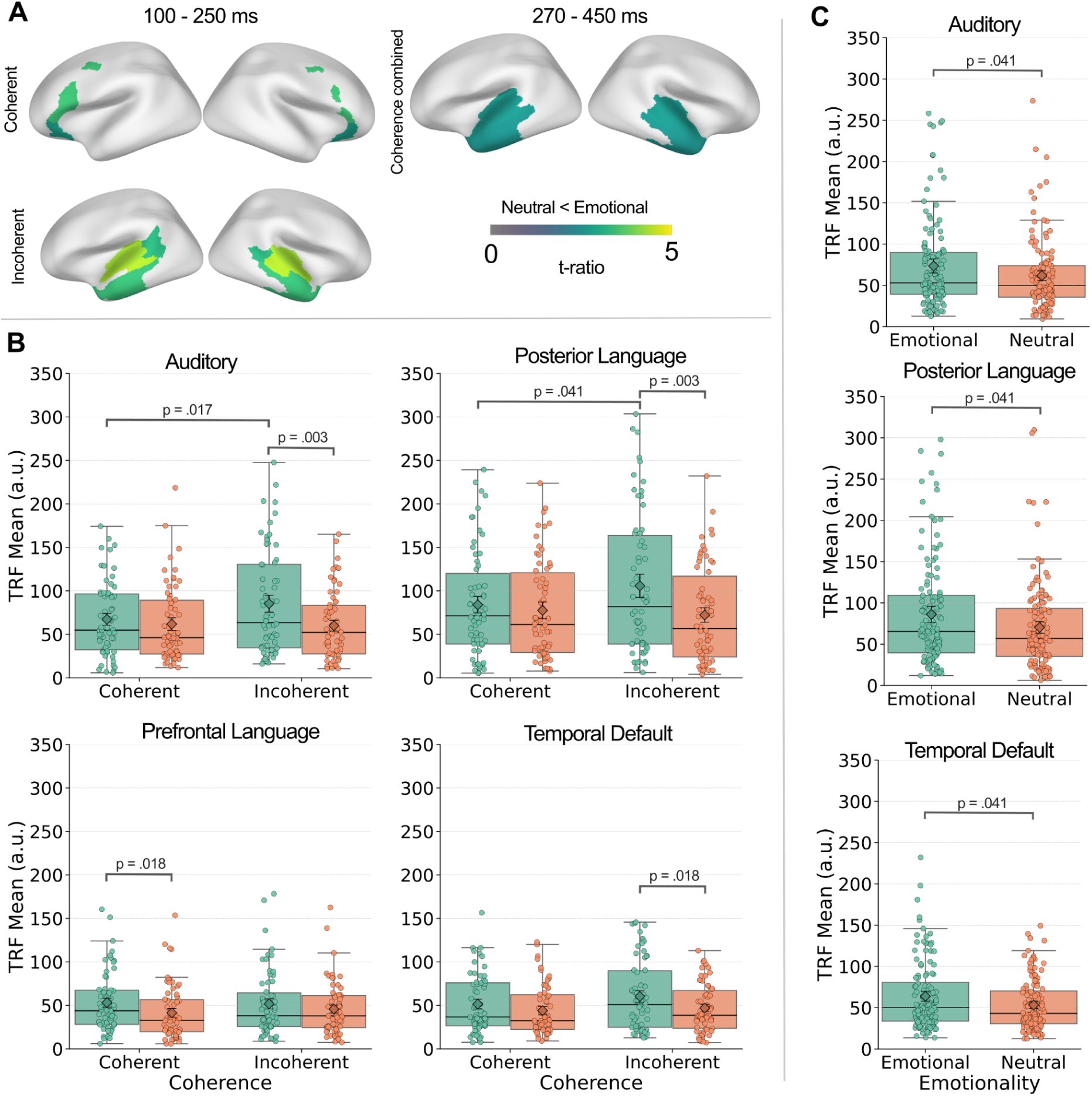
Emotional modulation of temporal response functions during attended speech. (A) Topographical distribution of significant differences (FDR-corrected) between emotional and neutral dialogues in the mid-latency (100–250 ms) and late-latency (270–450 ms) windows. Color intensity represents the strength of emotional enhancement (indexed by t-ratios). In the mid-latency window, coherence conditions are shown separately; in the late-latency window, coherence effects are not observed and conditions are collapsed. (B) Mid-latency TRF responses (100–250 ms) in representative ROIs for each condition combination. Boxplots show empirical distributions with individual observations overlaid (n = 31); diamonds indicate marginal means with standard errors. (C) Late-latency TRF responses (270–450 ms) in representative regions comparing emotional to neutral responses, collapsed across coherence conditions Boxplots show empirical distributions with individual observations overlaid (n = 31); diamonds indicate marginal means with standard errors.

In the late-latency window (270-450 ms), emotionally salient speech produced stronger responses than neutral speech (Fig. 4A, 4C; *F*(1, 30) = 7.56, *p* = .01, η*_G_*^2^ = 0.10), independent of dialogue coherence (p> .05, SI Table S9). Overall, late TRFs exhibited a broad enhancement for emotional speech in the temporal lobe that was less dependent on narrative structure than the mid-latency responses (SI Fig. S3). Together, these findings indicate that emotional meaning influences attentional tracking of conversations at two stages of speech comprehension: during initial derivation of lexico-semantic significance and later evaluative processing of socio-emotional meaning (Kotz & Paulmann, 2011; Schirmer & Kotz, 2006).

## Discussion

Using a combined encoding–decoding approach, we show that emotional semantic speech content robustly enhances sustained attentional engagement in conversation in naturalistic cocktail-party settings. By reconstructing the speech envelope from neural activity, we demonstrate that emotional dialogues enhance neural tracking of attended speech, with a distinct nonlinear temporal trajectory that is shaped by the semantic structure of the unfolding dialogue. Complementary Speech-to-brain encoding further reveals context-dependent and context-general effects of emotional salience across frontal, temporal and parietal brain regions. Together, these findings demonstrate that emotional salience of speech content shapes attention-related tracking of conversations by interacting with the turn-to-turn semantic context and engaging distributed cortical networks involved in auditory, semantic, and socio-cognitive processing.

Using listeners’ brain activity to reconstruct concurrent conversational speech, we found that emotional salience selectively enhanced neural tracking of attended, but not ignored, conversations. While enhanced tracking of attended speech is well established (Ding & Simon, 2012; Fiedler et al., 2019; O’Sullivan et al., 2015; Zion Golumbic et al., 2013), our results extend this work by showing that attentional tracking is further modulated by higher-order socio-emotional meaning. The absence of emotional effects for ignored speech indicates that emotional salience enhances attentional engagement only when speech is behaviorally relevant and subjected to full linguistic evaluation. This selectivity was mirrored in participants’ subjective ratings, as emotionally salient dialogues were consistently perceived as more affectively intense.

Tracking accuracy evolved nonlinearly across dialogue turns and depended on the dialogue’s semantic structure. Although prior work has shown that neural tracking fluctuates across conversational turns (Wikman et al., 2021, 2024), our findings demonstrate that emotional meaning shapes this dialogue-level evolution. In emotionally coherent dialogues, neural tracking showed a sharp early rise followed by attenuation, consistent with rapid stabilization of predictive models as emotional cues increase the precision of prediction errors (Schröger et al., 2015; Winkler et al., 2009). In contrast, emotionally incoherent dialogues exhibited a flatter trajectory with a later peak, consistent with sustained model updating and delayed convergence when salient input lacks coherent structure.

To link these dynamics to distinct neural processing stages, we complemented SER with TRF analyses, revealing effects in mid- and late-latency windows corresponding to established stages of emotional speech processing (Schirmer & Kotz, 2006; Kotz & Paulmann, 2011). At mid-latency, emotional salience enhanced responses in auditory, posterior language, and default mode regions during semantically incoherent dialogues, whereas coherent dialogues showed stronger effects in inferior frontal default mode and language regions. This pattern suggests that the earlier stages of emotional–semantic evaluation are sensitive to contextual predictability. At later latencies (270–450 ms), emotional speech elicited broader enhancements across temporal language and default mode regions regardless of coherence, consistent with sustained semantic and socio-cognitive integration. Together, these findings indicate that emotional meaning modulates speech processing at multiple stages through interactions between auditory–language networks and default mode regions supporting semantic comprehension and social cognition (Brodbeck et al., 2022; Citron, 2012; Fedorenko et al., 2024; Mar, 2011).

### Theoretical Implications

Taken together, these results support a hierarchical account in which emotional speech content sharpens attentional tracking at multiple levels of speech processing. At the lexical–semantic level, emotional meaning enhances tracking in a manner that depends on local semantic continuity between turns. At later stages of conceptual integration, emotional meaning enhances tracking independently of local continuity. Emotional salience thus modulates predictive processing across distinct hierarchical levels of the speech processing network.

Drawing from the expansion–renormalization framework of cortical plasticity (Kilgard, 2012), the results can be interpreted as emotionally salient content transiently increasing neural responsiveness through the recruitment of additional resources to meet task demands. In emotionally salient coherent dialogues, this increase is followed by a rapid renormalization as the neural system has adapted to task-relevant stimulus characteristics, returning to a less resource-intensive state. In contrast, semantic incoherence delays renormalization, resulting in slower adaptation and prolonged reliance on lower-level stimulus features.

Accordingly, emotional salience may transiently increase large-scale cortical organization, promoting coordinated engagement across perceptual, language, and control networks that supports effective response selection and action preparation (Clark, 2013; Diederen & Fletcher, 2021; Nau et al., 2021; Park et al., 2025; Song et al., 2021). In real-time conversation, such reorganization may prepare listeners for the initiation of a response, as conversational gaps are typically only 100–200 ms, requiring listeners to process incoming speech while concurrently preparing their own response (Brehm & Meyer, 2021; Bögels et al., 2015; Heldner & Edlund, 2010; Magyari, 2022; Stivers et al., 2009). Emotional salience may therefore facilitate the rapid turn-taking characteristic of emotionally engaging conversations (Stevanovic & Peräkylä, 2015).

### Limitations and future directions

In the present study, the emotional expressivity was limited to the semantic content of the dialogues, allowing us to explicitly study how content of speech modulates neural response associated with attentional engagement. However, human expressivity depends on the integration of various expressive modalities, such as paralinguistic features like changes in intonations, as well as communicative gestures like hand movements that likely co-occur with effects presented here. Future work should therefore extend the inquiry to conversations where the participants actively take part in the dialogue and are more free to use their communicative repertoire as a whole.

### Conclusions

Our findings show that emotional salience dynamically shapes attentional engagement in conversational speech in multi-speaker environments. Emotional semantic content enhances neural tracking and modulates the effect of contextual uncertainty on cortical responses. This interaction between emotion and predictability supports the view that auditory–language networks are adaptively tuned to socially and motivationally significant cues. Rather than acting as a simple amplifier of sensory encoding, emotional salience actively recruits and redistributes neural resources, reshaping language processing to meet changing conversational demands and enabling flexible adjustment between stable comprehension and adaptive responsiveness.

## Materials and Methods

### Participants

The sample comprised 32 participants (18 women, 14 men), aged between 20 and 50 years (*M* = 30.22, *SD* = 7.98) recruited through university mailing lists and campus outreach. The participants reported no neurological and psychiatric diagnoses or use of medications affecting the central nervous system, and had normal or corrected-to-normal vision and hearing. Before the experiment, all participants provided written informed consent. As compensation, participants received a reward valued at approximately 30 euros. The research protocol received approval from the University of Helsinki Research Ethics Committee of the Faculty of Medicine, and the study followed the ethical principles outlined in the World Medical Association’s Declaration of Helsinki. As one participant showed no variance in the speech-tracking signal during the processing phase, the dataset was excluded from further analyses.

### Experimental design

#### Procedure Overview

The participants completed an audiovisual “cocktail-party” experiment during MEG (Fig. 1A). The experiment, beginning after a 4-minute resting-state recording, consisted of two runs, with 16 trials each. In each trial, two six-line dialogues were presented simultaneously to the participants: an audiovisual target dialogue and an audio-only distractor dialogue, with audio presented from the same speakers concurrently. Participants were instructed to focus exclusively on the target dialogue.

To manipulate the semantic content in both target and distractor dialogues, the experiment followed a 4 * 4 factorial design, varying two dimensions of semantic content: the level of emotional salience (emotional vs. neutral) and the level of dialogue coherence at sentence level (narratively coherent dialogue vs. narratively incoherent dialogue). Emotional dialogues featured negatively valenced topics (e.g., tragic accident, fear- or disgust-related experience, conflict, health issue), whereas neutral dialogues described affectively non-salient activities (e.g., grocery shopping, attending a lecture, talking about the weather, drinking coffee). Coherent dialogues maintained contextually related conversational turns, while incoherent dialogues were generated by mixing lines from different dialogues. Each trial within a run formed a different factorial pairing, resulting in 16 unique target–distractor combinations. For example, both of the runs included a trial where an emotionally incoherent target dialogue was presented alongside a neutral coherent distractor dialogue.

#### Behavioral Ratings

To evaluate whether participants’ attended the target dialogues, they answered four true–false statements after each trial using a VPixx response device. The statements were designed to measure participants’ detection of central semantic information conveyed in the dialogues, such as recognizing mentioned people, places, objects, or actions (e.g., “The interactant asked whether the other person had an umbrella”; “The interactant is looking for a job”; “The interactant said that their lecturers change often”). Each true–false statement was constructed with its correct answer randomly set as either true or false. After responding, participants received feedback (e.g., “3 out of 4 correct”). Recall accuracy was computed as the proportion of correct responses (number of correct responses divided by four). Also, participants rated how emotionally charged the dialogue was on a three-point scale (1 = “Not at all”, 2 = “Somewhat”, 3 = “Highly”).

### Data Acquisition

#### Magnetoencephalography and Structural Magnetic Resonance Imaging

For source localization, structural MRI data were acquired using a 1.5 T Siemens Avanto scanner (Erlangen, Germany) with a T1-weighted 3D magnetization-prepared rapid gradient-echo (MP-RAGE) sequence (voxel size = 1 × 1 × 1 mm³). Brain activity during the experiment was recorded at 1 kHz with a 306-channel MEGIN Triux system (204 planar gradiometers, 102 magnetometers) at the BioMag Laboratory, Helsinki University Hospital. Horizontal and vertical electro-oculograms (EOG), as well as electrocardiograms (ECG), were recorded to detect eye movements and cardiac artifacts. The head position of the participants were continuously measured using 5 head position indicator (HPI) coils placed around the scalp. To facilitate accurate co-registration between the MEG data and a structural magnetic resonance image (MRI), 100 evenly divided scalp points were digitized (Polhemus).

### Data Processing

A detailed description of all data processing and analysis steps is provided in the Supporting Information (SI). The main procedures are summarized below.

#### MEG Preprocessing

MEG signals were first preprocessed using MNE-Python 1.5 (Gramfort et al, 2013). Raw recordings were denoised using MNE MaxFilter (Taulu & Simola, 2006) with temporal signal space separation (tSSS, 20-second sliding window). Data were high-pass filtered at 0.5 Hz and low-pass filtered at 15 Hz using zero-phase FIR filters. Artefacts were removed via the MNE FastICA -algorithm, and data were epoched based on dialogue turns for both target and distractor dialogues separately. Finally, the data were grouped based on emotional salience and semantic coherence, yielding separate neural data and audio envelope epochs for each Attention-Emotional Salience-Coherence combination.

#### Distributed Source Localisation

For estimating the cortical generators of the measured MEG signal, distributed minimum-norm source reconstruction (Gramfort et al., 2014; Hämäläinen & Ilmoniemi, 1994) was performed with each participant’s Freesurfer reconstructed surfaces. Single-shell boundary element head Model (BEM, conductivity 0.3 S/m) was computed from the inner skull surface, and minimum-norm inverse operators were built with dipole orientation constrained to the cortical surface normal using a loose orientation constraint. MEG data were projected to source space separately for each MEG epoch.

#### Stimulus processing

The broadband amplitude envelope of the dialogues were obtained as the absolute value of the Hilbert transform. Audio envelopes were band-pass filtered (0.5–15 Hz, 4th-order Butterworth filter), and sampling rate was matched to MEG data at 128 Hz. Finally, the envelopes were segmented into speech epochs for each line by detecting silent gaps of ≥1 s where the envelope stayed below 1% of the maximum amplitude. As the Distractor dialogue lines started 250 ms prior to the targets, the start of distractor epochs was cut to align with the onset of target lines.

### Encoding and Decoding Models

Decoding and encoding modelling were used to examine how distributed cortical activity tracked the auditory envelopes of the presented dialogues. By using speech envelope reconstruction (SER), it is possible to assess the fidelity at which the cortical activity follows the speech signal, whereas temporal response functions (TRFs) characterize the temporo-spatial dynamics of stimulus-response relationship (Crosse et al., 2016). The temporal lags considered in the models between speech signal and brain activity were restricted to be between -200 – 600 milliseconds. For both approaches, the *receptive field estimation* function in MNE-Python was used, with the default lagged ridge-regression approach, with 4-fold crossvalidation (regularisation-parameters tested between 1 to 1 * 10^6^; see SI Fig. S4). Separate models were fit for each Attention - Emotional Salience - Dialogue coherence combination, and model fits were assessed based on the fit of each condition, ultimately providing the best fit was regularization at 1 * 10^4^ for TRFs. Same regularisation was chosen for SER.

#### Speech Envelope Reconstruction (SER)

To reduce computational strain, the neural input data was first condensed to anatomically-functionally defined cortical areas, based on the human connectome project’s multi-modal parcellation (Glasser et al., 2016), by calculating the average time-course within each cortical area (with sign-flip of activations in the non-dominant orientations with > 90 degree deflection). Reconstruction accuracy was quantified as Pearson’s product-moment correlation coefficient of the reconstructed and true envelopes across cross-validation folds. To assess the temporal evolution of neural speech representations, the cross-validation test reconstructions from the final chosen model were back-projected to their original temporal locations to build the full reconstructed speech envelope. Then, the mean correlation between SER and the stimulus envelope was calculated for each turn. The overall mean response and responses varying across turns were subjected to separate statistical analysis, with the correlation coefficients Fisher’s *z*-transformed prior to analysis.

#### Temporal Response Functions (TRFs)

TRFs were used to model how changes in the speech envelope drove cortical responses over time with the envelope as the input and the neural source activity at each vertex as the response. Regression coefficients were averaged across folds to yield source-by-time TRFs for each subject and condition and were subsequently averaged based on the glasser cortical parcellation. From this partition, the auditory, language and default-mode related regions located in the lateral temporal, parietal and frontal areas were chosen for further analysis. As the grand averaged responses displayed a distinct two-peak response pattern, the data were analysed in two separate ime windows of interest. For the earlier window, a local positive peak within 100–250 ms was identified for each network and condition separately. The mean amplitude within a 50 ms window around this peak was computed, capturing the mid-latency response while accommodating latency differences across cortical regions. The later activation was quantified as the overall mean across 270-450 ms time window, due to less pronounced curvature in the source waveforms.

### Statistical Analyses

All statistical analyses were conducted in R (version 4.5.1). Type-III repeated measures analysis of variances (rm-ANOVAs) were done with the afex package. Post hoc comparisons were based on estimated marginal means (emmeans package) with Holm correction for multiple comparisons in SER, and FDR-correction in TRF-analysis. For perceived emotional load, we used generalized estimating equation (GEE) for clustered ordinal responses (ordgee, geepack).

## Acknowledgements

The authors would like to thank Mariel Wuolio for her valuable help in preparing materials, participant recruitment and assistance in MEG measurements. We are also grateful to Markus Söderman for his assistance with the preparation of the stimulus material and study design, and Jan Lindström for his role in the conceptualization and planning of the project. Finally, we thank students Leevi Lintunen, Jenna Jakonen, Aino Innanen and Aarni Arvilommi for their valuable contributions in producing the study’s stimulus material.

