## Supplementary material for "Emotional Content Enhances Neural Tracking of Conversations": SI

### Supporting Information for
Emotional Content Enhances Neural Tracking of Conversations

Heikkinen, P.[1,4]*, Lepistö, S.[1]*, Wikman, P.[1,2,3,4], Koskinen, E.[5], Peräkylä, A.[5], Ravaja, N.[1], & Harjunen, V. J.[1]

*Shared first

[1] *Department of Psychology, Faculty of Medicine, University of Helsinki, Helsinki, Finland*

[2] *Department of Neuroscience, Georgetown University, Washington D.C., USA*

[3] *Advanced Magnetic Imaging Centre, Aalto NeuroImaging, Aalto University, Espoo, Finland*

[4] *Finnish Centre of Excellence in Music, Mind, Body, and Brain, Cognitive Brain Research Unit, Department of Psychology, Faculty of Medicine, University of Helsinki, Helsinki, Finland*

[5] *Faculty of Social Sciences, University of Helsinki, Helsinki, Finland*

**Corresponding Author**: Pyry Heikkinen

**This PDF file includes:**

Supporting text
Figures S1 to S4
Tables S1 to S9
SI References

**Supporting Information Text**
**SI Methods**

**Dialogue Material.** The stimulus material for 32 target and 32 distractor dialogues presented in each measurement were drawn from a larger dialogue repository containing a total of 128 different dialogues, varying across the two dimensions of emotional salience (emotional vs. neutral) and coherence (coherent vs. incoherent). All coherent emotional and neutral dialogues varied in their conversational topics (but matched for length and structure), while incoherent dialogues were created by mixing lines from the coherent dialogues (ensuring that no lines originating from the same original dialogue appeared together in the same incoherent version). Consequently, the level of semantic coherence was manipulated by varying semantic relatedness of the interactants' utterances: in coherent dialogues, interactants responded appropriately to each other's speech (e.g., one asks about the co-interactant's experience, and the other replies with information about that experience), whereas in incoherent dialogues, the turns were not semantically connected (e.g., one talks about gardening, the other talks about shopping). In turn, the level of emotional salience was manipulated by using dialogues that were either emotional (e.g., tragic accident, fear- or disgust-related experience, physical conflict, mental health issue or illness, harassment, loneliness) or neutral (e.g., grocery shopping, attending a lecture, cooking, working, talking about the weather, having lunch, drinking coffee), with all emotional dialogues being negatively valenced. Each dialogue was available in two versions, performed by different male–female pairs (Dyad A, Dyad B). These pairs were counterbalanced across participants: for half of the participants, Dyad A performed the target dialogues while Dyad B performed the distractors; for the other half, this assignment was reversed. The gender of the active speaker was kept different between the target and distractor conversations. All dialogues comprised a six-line exchange (lasting approximately 30 seconds in total), with each interactant contributing three lines. As participants also answered four true–false statements after each trial, four of the six lines were randomly selected (and a corresponding true–false statement was created for each selected line). Furthermore, as the study focused on the neural tracking of semantic content, we minimized the effects of other communicative features. Importantly, interactants of the dialogues did not exhibit bodily expressions (e.g., facial expressions, nodding, hand gestures, or postural movement). Crucially, they delivered speech with deliberately flat prosody, minimizing variation in stress, rhythm, and loudness across dialogue lines.

**Stimulus Presentation**. The stimulus was presented with the Presentation® software (Neurobehavioral Systems Inc.). The target dialogues were presented on a projected screen (67 * 38 cm) at a viewing distance of approximately 130 cm (from the participant's eyes). The projector's resolution was 1920 * 1080 (Full HD), with a refresh rate of 120 Hz. Before each target dialogue, a gray screen with the instruction "Pay attention to the conversation" was shown. Subsequently, target and distractor dialogues were presented line by line. After each line, a gray screen with a fixation cross was presented. For each line, the stimulus volume was set to a comfortable level based on the target, and the paired distractor was attenuated by 10 decibels based on the root mean square of the amplitudes, and each distractor line was set to begin 250 ms before the onset of the corresponding target line.

**MEG Preprocessing**. MEG data were preprocessed using MNE-Python 1.5 (Gramfort et al, 2013). Raw recordings were first denoised using MNE MaxFilter (Taulu & Simola, 2006) with temporal signal space separation (tSSS, 20-second sliding window, with the system's calibration and crosstalk files). The head position of each participant was spatially aligned to the start of run 1, with movement compensation based on the recorded continuous HPI coil information. Noisy and flat channels were automatically marked  and the data was resampled at 128 Hz for processing efficiency. Data were high‑pass filtered at 0.5 Hz and low‑pass filtered at 15 Hz using zero‑phase FIR filters. Bad channel selection was manually confirmed, and channels were interpolated with spherical‑spline interpolation. Next, ocular and cardiac artefacts were removed via the MNE FastICA -algorithm by flagging components correlated EOG and ECG, respectively, and manually confirming the ICA exclusion. Next, MEG data were epoched from each dialogue line onset to offset for both target and distractor dialogues separately. Finally, the data were grouped based on emotional salience and semantic coherence, yielding separate neural data and audio envelope epochs for each Attention-Emotional Salience-Coherence combination.

**Distributed Source Localisation**. For estimating the cortical generators of the measured MEG signal, distributed minimum-norm source reconstruction (Gramfort et al., 2014; Hämäläinen &

Ilmoniemi, 1994) was performed with each participant's Freesurfer reconstructed surfaces. The surfaces were coordinate aligned to MEG digitization points, and a single-shell boundary element head Model (BEM, conductivity 0.3 S/m) was computed from the inner skull surface. The cortical source spaces were defined at oct6 resolution (~5 mm spacing) on the white matter surface. Noise covariance matrices were estimated from 4 minute long empty-room recordings. Minimum-norm inverse operators were built (depth weighting = 0.2; SNR = 1/9) with dipole orientation constrained to the cortical surface normal using a loose orientation constraint (0.2). MEG data were projected to source space separately for each MEG epoch, yielding one cortical source current estimate time-locked to each stimulus combination.

#### Figures

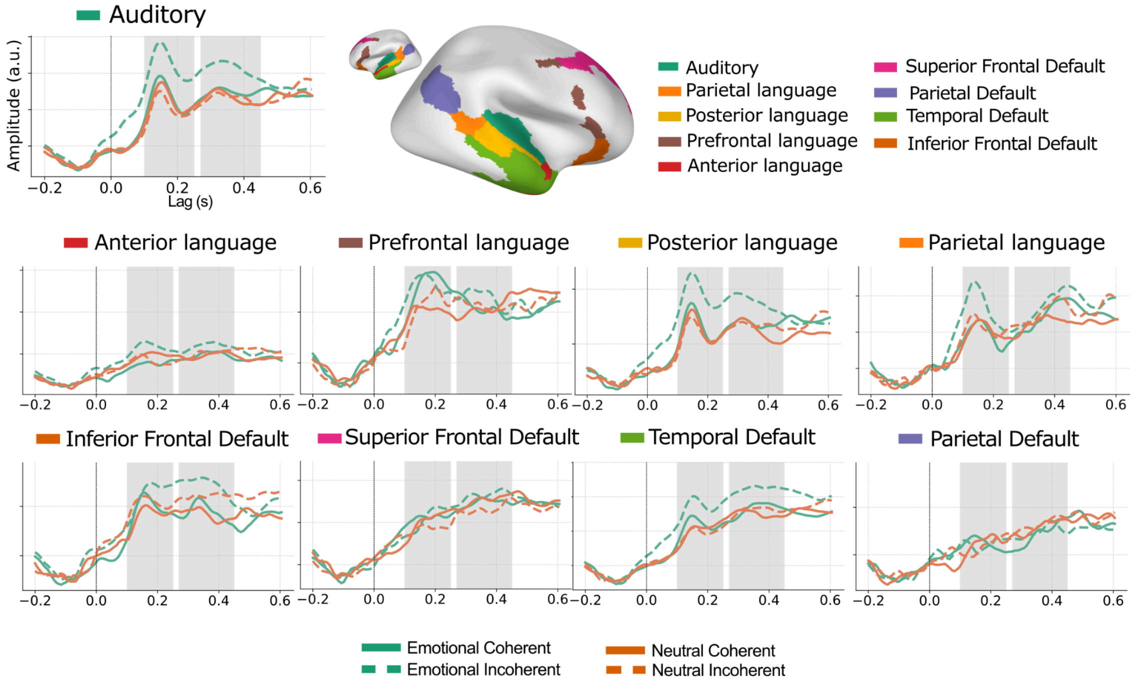


**Fig. S1.** Right-hemisphere regions of interest used for TRF analyses. Cortical parcels in the right hemisphere corresponding to auditory, language, and default-mode network regions included in the TRF analyses. Regions mirror those shown for the left hemisphere in the main text and were selected based on prior work on speech and narrative comprehension. Note that hemisphere specific effects were not analyzed.

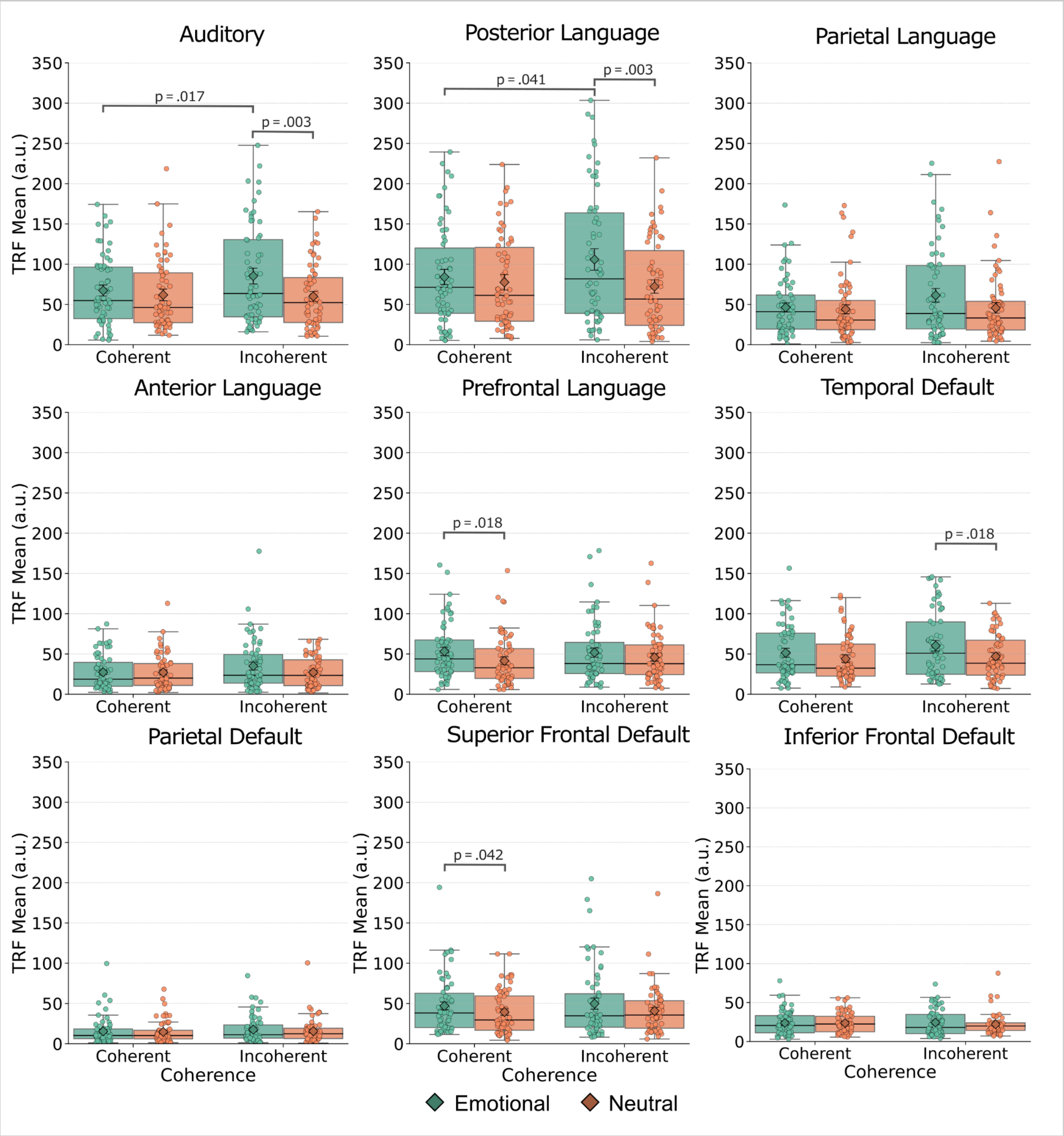


**Fig. S2.** Mid-latency TRF responses across all regions of interest. Mean TRF amplitudes (100–250 ms) for attended speech are shown for each region of interest and condition combination (emotional vs. neutral; coherent vs. incoherent). Boxplots depict empirical distributions across participants (n = 31), with individual observations overlaid. Diamonds indicate estimated marginal means with standard errors.

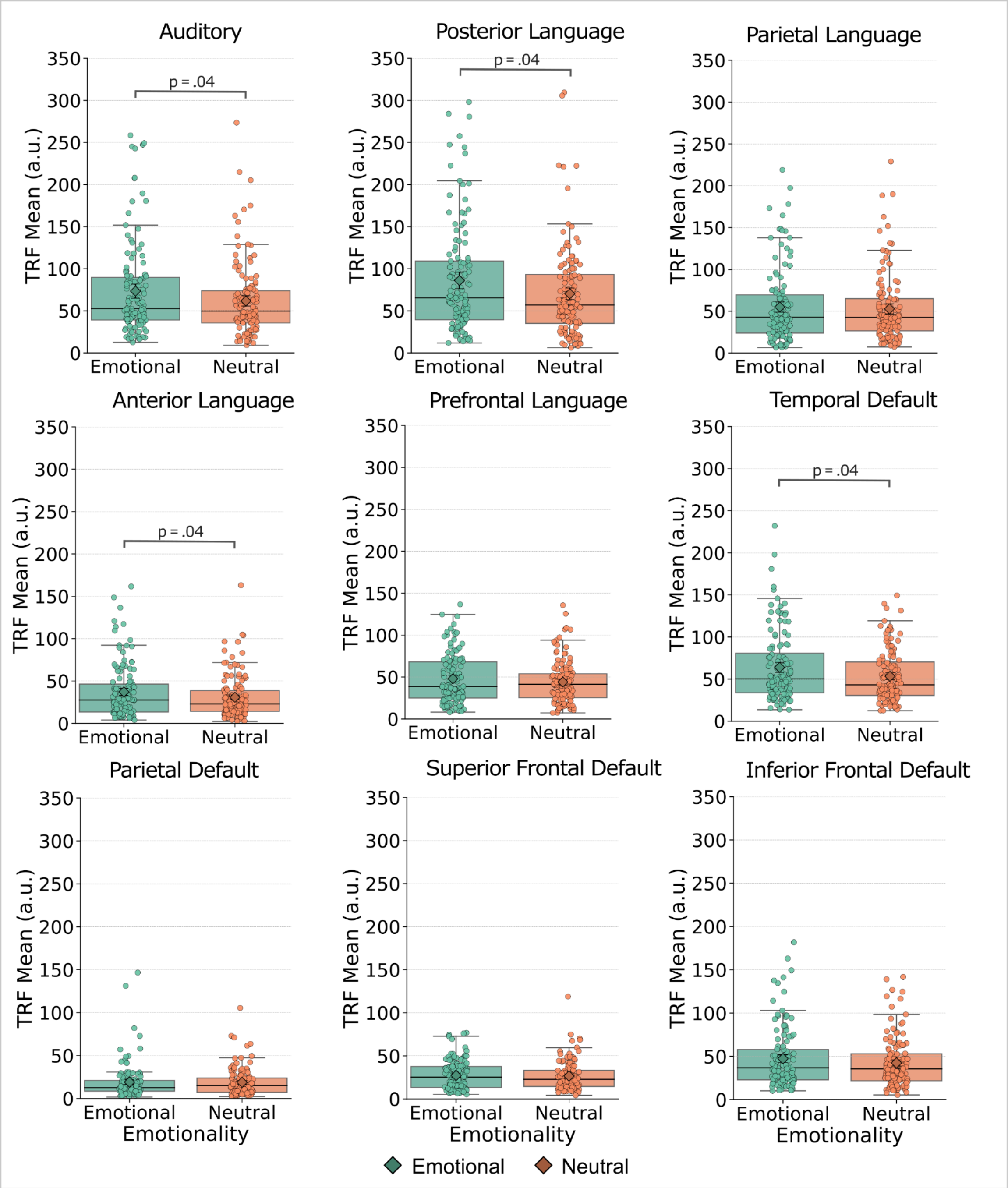


**Fig. S3.** Late-latency TRF responses across all regions of interest. Mean TRF amplitudes (270–450 ms) for attended speech are shown for each region of interest, collapsed across dialogue coherence. Boxplots show empirical distributions across participants (n = 31), with individual observations overlaid. Diamonds indicate linear mixed-effects model marginal means with standard errors.

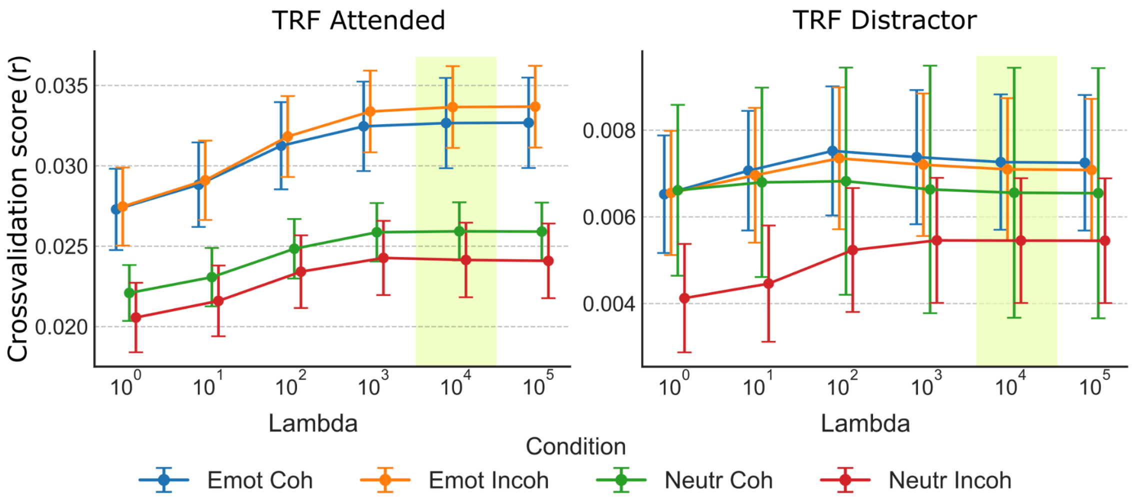


**Fig. S4.** Cross-validated TRF model performance for each emotional salience × dialogue coherence condition in targer and distractor trials. Crossvalidation score was quantified as the Pearson correlation between predicted and observed neural responses using four-fold cross-validation. The shaded vertical band indicates the regularization parameter (λ) selected for all subsequent TRF analyses.

**Tables**

**Table S1.** Results of an ordinal generalized estimating equation (GEE) model testing the effects of emotional salience (emotional vs. neutral), dialogue coherence (coherent vs. incoherent), and their interaction on perceived emotional load ratings. Wald $\chi^2$ statistics are reported for each predictor.

| Emotional Load: Ordinal GEE | | | | |
|---|---|---|---|---|
| Effect | Estimate | SE | Wald | p |
| emotionality | 14.144 | 0.136 | 10,867.51 | < .001 |
| coherence | 0.075 | 0.040 | 3.48 | = 0.062 |
| emotionality × coherence | -0.003 | 0.038 | 0.01 | = 0.930 |

**Table S2.** Recall accuracy repeated-measures ANOVA. Type-III repeated-measures ANOVA testing the effects of emotional salience (emotional vs. neutral), and dialogue coherence (coherent vs. incoherent) on recall accuracy. Generalized eta-squared ($\eta^2$) is reported as an effect size.

| Recall Accuracy: Type-III ANOVA | | | | | |
|---|---|---|---|---|---|
| Effect | df1 | df2 | F | ges | p |
| emotionality | 1 | 30 | 0.81 | 0.004 | = 0.376 |
| coherence | 1 | 30 | 53.73 | 0.199 | < .001 |
| emotionality × coherence | 1 | 30 | 0.08 | 0.000 | = 0.779 |

**Table S3.** SER factorial repeated-measures ANOVA. Type-III repeated-measures ANOVA testing the effects of attention (attended vs. distractor), emotional salience (emotional vs. neutral), and dialogue coherence (coherent vs. incoherent) on speech envelope reconstruction (SER) accuracy. Generalized eta-squared ($\eta^2$) is reported as an effect size.

| SER by factor: Type-III ANOVA | | | | | |
|---|---|---|---|---|---|
| Effect | num Df | den Df | F | etasquared | p |
| attention | 1 | 30 | 408.79 | 0.56 | < .001 |
| emotionality | 1 | 30 | 26.66 | 0.06 | < .001 |
| coherence | 1 | 30 | 0.05 | 0.00 | = .831 |
| attention:emotionality | 1 | 30 | 13.11 | 0.05 | = .001 |
| attention:coherence | 1 | 30 | 0.36 | 0.00 | = .555 |
| emotionality:coherence | 1 | 30 | 2.31 | 0.01 | = .139 |
| attention:emotionality:coherence | 1 | 30 | 6.17 | 0.02 | = .019 |

**Table S4.**SER post-hoc pairwise comparisons. Holm-adjusted pairwise comparisons of estimated marginal means (EMMs) for SER accuracy, conducted separately for attended and distractor speech across emotional salience and dialogue coherence conditions. Significant contrasts demonstrate that emotional salience enhanced SER selectively for attended speech.

| Pairwise contrasts (Holm-adjusted) for SER EMMs | | | | | | | | |
|---|---|---|---|---|---|---|---|---|
| attention | contrast | estimate | SE | df | t.ratio | 95% -confidence | upper.CL | p |
| trgt | emo coh - neut coh | 0.047 | 0.008 | 30 | 5.933 | 0.025 | 0.069 | < .001 |
| | emo coh - emo incoh | 0.000 | 0.010 | 30 | -0.015 | -0.028 | 0.028 | = 0.988 |
| | emo coh - neut inco | 0.038 | 0.013 | 30 | 2.934 | 0.001 | 0.074 | = 0.019 |
| | neut coh - emo incoh | -0.047 | 0.010 | 30 | -4.651 | -0.076 | -0.018 | < .001 |
| | neut coh - neut incoh | -0.009 | 0.012 | 30 | -0.742 | -0.044 | 0.026 | = 0.928 |
| | emo incoh - neut incoh | 0.038 | 0.012 | 30 | 3.135 | 0.004 | 0.072 | = 0.015 |
| dist | emo coh - neut coh | -0.019 | 0.010 | 30 | -1.912 | -0.047 | 0.009 | = 0.262 |
| | emo coh - emo incoh | -0.020 | 0.013 | 30 | -1.497 | -0.058 | 0.018 | = 0.435 |
| | emo coh - neut inco | 0.005 | 0.009 | 30 | 0.514 | -0.020 | 0.029 | = 1.000 |
| | neut coh - emo incoh | -0.001 | 0.012 | 30 | -0.089 | -0.036 | 0.034 | = 1.000 |
| | neut coh - neut incoh | 0.023 | 0.011 | 30 | 2.223 | -0.006 | 0.053 | = 0.203 |
| | emo incoh - neut incoh | 0.024 | 0.011 | 30 | 2.158 | -0.008 | 0.056 | = 0.203 |

**Table S5.** Line-wise repeated-measures ANOVA for SER accuracy. Type-III repeated-measures ANOVA examining the effects of conversational turn (line), emotional salience, and dialogue coherence on SER accuracy for attended speech. Results reveal a significant three-way interaction, indicating that emotional salience and coherence jointly shape the temporal evolution of neural speech tracking across dialogue turns.

| Targets: Type-III RM-ANOVA (line × emotionality × coherence) | | | | | |
|---|---|---|---|---|---|
| Effect | num DF | den DF | F | etasquared | p |
| emotionality | 1.00 | 30.00 | 31.04 | 0.06 | < .001 |
| coherence | 1.00 | 30.00 | 0.21 | 0.00 | = .652 |
| line | 4.02 | 120.56 | 8.47 | 0.04 | < .001 |
| emotionality:coherence | 1.00 | 30.00 | 0.24 | 0.00 | = .627 |
| emotionality:line | 4.32 | 129.71 | 3.18 | 0.02 | = .013 |
| coherence:line | 4.33 | 129.92 | 3.33 | 0.02 | = .010 |
| emotionality:coherence:line | 4.23 | 126.98 | 4.58 | 0.03 | = .001 |

**Table S6.** Polynomial contrast comparisons between emotional and neutral dialogues. Holm-adjusted pairwise comparisons of linear, quadratic, and cubic polynomial contrasts comparing emotional and neutral dialogues, conducted separately for coherent and incoherent conditions. Significant quadratic contrasts indicate stronger nonlinear trajectories for emotional speech, particularly in coherent dialogues.

| contrast | coherence | estimate | SE | df | t.ratio | lower.CL | upper.CL | p |
|---|---|---|---|---|---|---|---|---|
| linear | coherent | -0.032 | 0.192 | 30 | -0.169 | -0.424 | 0.360 | = 0.867 |
| quadratic | | -0.596 | 0.157 | 30 | -3.797 | -0.916 | -0.275 | < .001 |
| cubic | | 0.276 | 0.274 | 30 | 1.008 | -0.283 | 0.836 | = 0.322 |
| linear | incoherent | 0.130 | 0.175 | 30 | 0.745 | -0.227 | 0.488 | = 0.462 |
| quadratic | | -0.199 | 0.174 | 30 | -1.145 | -0.553 | 0.156 | = 0.261 |
| cubic | | -0.382 | 0.260 | 30 | -1.469 | -0.913 | 0.149 | = 0.152 |

**Table S7.** Polynomial contrast comparisons between coherent and incoherent dialogues. Holm-adjusted pairwise comparisons of linear, quadratic, and cubic polynomial contrasts comparing coherent and incoherent dialogues, conducted separately for emotional and neutral conditions. Significant cubic contrasts for emotional dialogues indicate coherence-dependent shifts in the timing of peak SER accuracy.

| contrast | emotionality | estimate | SE | df | t.ratio | lower.CL | upper.CL | p |
|---|---|---|---|---|---|---|---|---|
| linear | emotional | -0.173 | 0.148 | 30 | -1.170 | -0.476 | 0.129 | = 0.251 |
| quadratic | | -0.264 | 0.168 | 30 | -1.570 | -0.607 | 0.079 | = 0.127 |
| cubic | | 0.874 | 0.284 | 30 | 3.075 | 0.293 | 1.454 | = 0.004 |
| linear | neutral | -0.011 | 0.188 | 30 | -0.056 | -0.394 | 0.373 | = 0.955 |
| quadratic | | 0.133 | 0.165 | 30 | 0.806 | -0.204 | 0.470 | = 0.427 |
| cubic | | 0.216 | 0.272 | 30 | 0.791 | -0.341 | 0.772 | = 0.435 |

**Table S8.** Mid-latency TRF repeated-measures ANOVA. Type-III repeated-measures ANOVA testing the effects of emotional salience, dialogue coherence, cortical roi, and hemisphere on mid-latency (100–250 ms) TRF amplitudes for attended speech. Generalized eta-squared (ges) is reported as an effect size. Results reveal a significant Emotion × Coherence × roi interaction, indicating context-dependent modulation of early emotional–semantic processing across auditory, language, and default-mode network regions.

| Effect | df1 | df2 | F | ges | p |
|---|---|---|---|---|---|
| emotionality | 1.00 | 30.00 | 17.17 | 0.014 | < .001 |
| coherence | 1.00 | 30.00 | 3.43 | 0.003 | = .074 |
| roi | 2.16 | 64.79 | 44.02 | 0.234 | < .001 |
| hemisphere | 1.00 | 30.00 | 1.68 | 0.002 | = .205 |
| emotionality × coherence | 1.00 | 30.00 | 6.22 | 0.003 | = .018 |
| emotionality × roi | 3.50 | 105.09 | 5.73 | 0.006 | < .001 |
| coherence × roi | 3.14 | 94.24 | 1.46 | 0.002 | = .229 |
| emotionality × hemisphere | 1.00 | 30.00 | 0.00 | 0.000 | = .949 |
| coherence × hemisphere | 1.00 | 30.00 | 0.55 | 0.000 | = .463 |
| roi × hemisphere | 3.27 | 98.12 | 2.67 | 0.008 | = .047 |
| emotionality × coherence × roi | 4.11 | 123.21 | 4.96 | 0.004 | < .001 |
| emotionality × coherence × hemisphere | 1.00 | 30.00 | 1.30 | 0.000 | = .263 |
| emotionality × roi × hemisphere | 3.52 | 105.52 | 1.23 | 0.001 | = .302 |
| coherence × roi × hemisphere | 3.25 | 97.54 | 0.32 | 0.000 | = .826 |
| emotionality × coherence × roi × hemisphere | 3.77 | 112.98 | 0.84 | 0.001 | = .497 |

**Table S9** .Late-latency TRF repeated-measures ANOVA. Type-III repeated-measures ANOVA testing the effects of emotional salience, dialogue coherence, cortical roi, and hemisphere on late-latency (270–450 ms) TRF amplitudes for attended speech. Generalized eta-squared (ges) is reported as an effect size. Results show a robust main effect of emotional salience and region-specific modulation, largely independent of dialogue coherence.

| Effect | df1 | df2 | F | ges | p |
|---|---|---|---|---|---|
| emotionality | 1.00 | 30.00 | 7.56 | 0.007 | = .010 |
| coherence | 1.00 | 30.00 | 1.25 | 0.001 | = .273 |
| roi | 2.37 | 71.09 | 34.91 | 0.197 | < .001 |
| hemisphere | 1.00 | 30.00 | 9.07 | 0.017 | = .005 |
| emotionality × coherence | 1.00 | 30.00 | 0.51 | 0.000 | = .479 |
| emotionality × roi | 3.04 | 91.19 | 3.60 | 0.005 | = .016 |
| coherence × roi | 3.22 | 96.71 | 0.48 | 0.001 | = .708 |
| emotionality × hemisphere | 1.00 | 30.00 | 1.62 | 0.000 | = .213 |
| coherence × hemisphere | 1.00 | 30.00 | 3.45 | 0.002 | = .073 |
| roi × hemisphere | 3.46 | 103.82 | 2.22 | 0.008 | = .081 |
| emotionality × coherence × roi | 4.20 | 125.87 | 0.90 | 0.001 | = .468 |
| emotionality × coherence × hemisphere | 1.00 | 30.00 | 1.25 | 0.000 | = .272 |
| emotionality × roi × hemisphere | 4.68 | 140.25 | 0.82 | 0.001 | = .530 |
| coherence × roi × hemisphere | 3.97 | 119.18 | 2.14 | 0.002 | = .080 |
| emotionality × coherence × roi × hemisphere | 2.91 | 87.29 | 1.04 | 0.001 | = .376 |
